# Costs of phenotypic plasticity can undermine its benefits for evolvable digital organisms

**DOI:** 10.1101/2023.03.13.532337

**Authors:** Karine Miras

**Affiliations:** Department of Computer Science, Vrije Universiteit Amsterdam, Netherlands

## Abstract

Phenotypic plasticity is usually defined as a property of individual genotypes to produce different phenotypes when exposed to different environmental conditions. While the benefits of plasticity for adaptation are well established, the costs associated with plasticity remain somewhat obscure. Understanding both why and how these costs occur could help us explain and predict the behaviour of living creatures as well as allow us to design more adaptable robotic systems. One of the challenges of conducting such investigations concerns the difficulty in isolating the effects of different types of costs and the lack of control over environmental conditions. The present study tackles these challenges by using virtual worlds (software) to investigate the environmentally regulated phenotypic plasticity of digital organisms: the experimental setup guarantees that possibly incurred genetic costs of plasticity are isolated from other plasticity-related costs. The hypothesis put forward here is that despite the potential benefits of plasticity, these benefits might be undermined by the genetic costs related to plasticity itself. This hypothesis was subsequently confirmed to be true.

**Author summary:** Phenotypic plasticity is usually defined as a property of individual DNA that produces different bodies and brains when exposed to different environmental conditions. While the benefits of plasticity for adaptation are well established, there are also potential costs associated with plasticity: “Jack of all trades, master of none.” Understanding both why and how these costs occur could help us explain and predict the behaviour of living creatures as well as allow us to design more adaptable robotic systems. While some studies have reported strong evidence for such costs, many other studies have observed no costs. One of the challenges associated with conducting such investigations concerns the difficulty of isolating the effects of the different types of costs. Artificial life (ALife) involves the design and investigation of artificial living systems in different levels of organisation and mediums. Importantly, ALife allows for the customisation of multiple properties of an artificial living system. In the present study, I investigate the environmentally regulated phenotypic plasticity of evolvable digital organisms using an ALife system. The experimental setup guarantees that possibly incurred genetic costs of plasticity are isolated from other plasticity-related costs. The hypothesis put forward here is that despite the potential benefits of plasticity, these benefits might be undermined by the genetic costs related to plasticity itself. This hypothesis was subsequently confirmed to be true.

## Introduction

In nature, the traits of living creatures are not coded independently in their genetic material. One obvious reason for this is compression: the reuse of genetic material helps to simplify this ‘optimization problem’ by solving small parts and then combining them accordingly. For instance, while only approximately 30, 000 genes code all traits of the human phenotype [1], the human brain alone consists of trillions of neurons [2]. While compression is expedient for simple cases of reuse, e.g., coding for multiple fingers, it also helps to produce more profound benefits: the remarkable levels of intelligence observed in nature are not simply a result of the discovery of how to produce certain traits, but rather, most importantly, derive from solving when to express these traits [3].

The environment surrounding organisms is constantly changing, and therefore organisms might encounter countless environmental conditions over their lifetime. These changes can vary from internal (intra-phenotype) to external changes, such as, for example, the weather, terrain, predators, inter-species interactions, etc. To cope with these changes, different dimensions of phenotypic development are observed in nature.

Phenotypic plasticity “is usually defined as a property of individual genotypes to produce different phenotypes when exposed to different environmental conditions” [4]. The term encompasses morphological, physiological, and behavioural traits and has been utilised in a very broad sense throughout extant literature [5], including even within the fields of learning [6] and body training [7]. The more specific usage of the term refers to the environmentally regulated expressions of different phenotypes from the same genotype. One example of this is polyphenism [8] - non-reversible phenotypic differentiation, e.g., castes in social insects and sex determination in reptiles. Another example is acclimatization [9] - reversible phenotypic changes triggered by environmental cues, e.g., Passerine birds that change their musculature to cope with winter.

While the benefits of plasticity are well established, the constraints surrounding plasticity remain somewhat obscure [10]. Such constraints are referred to as *costs* and *limits*. Although these costs and limits have been divided into distinct categories in extant literature, their exact definitions diverge, and their mechanisms are not yet well understood [10–12]. One definition describes them as follows: “Costs of plasticity are fitness deficits associated with the plastic genotypes relative to fixed genotypes producing the same mean phenotype in a focal environment, and limits of plasticity are functional constraints that reduce the benefit of plasticity compared to perfect plasticity.” [10] This means that in the case of limits, the plastic genotype can not express the (extreme) traits like the non-plastic genotype does, while in the case of costs, the traits can be achieved, but the fitness is reduced for other reasons. Another definition argues that expressing sub-optimal (‘wrong’) phenotypes in a given environment could be viewed as a cost, and that it is not straightforward to separate costs from limits [13].

Though plasticity-related costs have been studied throughout the years, the evidence explaining both when and how these constraints occur remains insufficient [12]. Previous work has produced a mixed picture of the importance of plasticity-related costs: while some studies report strong evidence for costs, but many other studies observe no costs [14].

There are a wide range of challenges that make it difficult to study plasticity-related costs, which explains some of the disagreement amongst the studies. One of these challenges pertains to the potential differences between the environments where acclimation response evolved and the environments used for tests and analysis, while another difficulty concerns the lack of control for the types of plasticity [15]. Moreover, there are multiple types of costs, which could hardly be experimentally isolated in natural life systems. Five classical costs have been defined in the literature: *production costs, maintenance costs, information-acquisition costs, developmental-instability costs*, and *genetic costs* [10,13].

The present study tackles these challenges through Artificial life (ALife). ALife has been characterised as the study of “life as it could be” [16], and it involves the design and investigation of artificial living systems in different levels of organisation (molecular, cellular, organismal, populational) and mediums (hardware, software, wetware) [17]. Advances in ALife are expedient for designing artificially intelligent systems as well as for understanding naturally intelligent systems. There are at least three benefits of using ALife to investigate natural phenomena. First, it allows for the customisation of multiple properties within an artificial living system. Evidently, this does not guarantee the emergence of specific phenomena, but rather makes it possible to deliberately include or exclude certain mechanisms that may (or may not) lead to the emergence of these phenomena. This flexibility is relevant for studying phenotypic plasticity costs because it makes it possible to isolate effects within different types of costs. Second, it enables control over environmental conditions. Finally, it allows for the relatively cheap repeatability of experiments for statistical significance.

Artificial organisms that possess both a body and a brain have been evolved in several studies since 1994 [18–20]. However, scarce attention has been paid to the environmental regulation of phenotypic plasticity. Whereas only a few examples of studies related to development and phenotypic plasticity are found in the literature [21–27], none of these examine the costs of plasticity. Here, a hypothesis is posited that despite the potential of phenotypic plasticity, its benefits may be potentially undermined by plasticity-related genetic costs.

The current experimental setup evolves multiple populations of organisms endowed with and without phenotypic plasticity in simulation. The resulting experiments are used to investigate the *genetic costs* of phenotypic plasticity (specifically acclimatisation) upon both the behaviour and phenotypic traits of evolvable digital organisms. Genetic costs are those “costs that result directly from linkage between loci affecting plasticity and loci with negative fitness effects, (negative) pleiotropic effects of loci affecting plasticity and other traits, or (negative) epistatic interactions among loci affecting plasticity and other loci [13].” Genetic linkage [28] refers to the tendency of genetic sequences located nearby on a chromosome to be inherited together during sexual reproduction. *Pleiotropy* [29] concerns genes that are able to influence more than one trait in the phenotype. *Epistasis* [30] refers to the interaction between genes, namely the fact that the effect of a gene may depend on either the presence or absence of another gene(s). Both pleiotropy and epistasis are possible within the utilised genetic representation. Linkage does not apply because reproduction within the current system is asexual.

Given that this work is aimed at audiences from various backgrounds and, as such, there can sometimes be a terminology mismatch, the key terms used throughout the text are clarified in S1 Appendix.

## Experimental Setup

All experiments (each setup) described here were repeated on *20* occasions independently for statistical significance. The experiments were carried out using six different setups that combined three aspects: phenotypic plasticity (with or without), phenotypic component (body or brain), and desired behavior (locomote left, locomote right, locomote down). In these experiments, the organisms had to perform a pair of behaviours throughout the 60 seconds of their lifetime: in a plane terrain environment, they were supposed to locomote to direction A in the first 30 seconds of their life and then locomote to direction B in the last 30 seconds of their life. To ensure that the behavioural metrics were as comparable as possible, the organisms were reset to their starting position before moving on to direction B. In this way, for both directions A and B, the initial conditions of the body morphology were the same.

The pairs of directions were *right-left* and *right-down*. Methods with and without phenotypic plasticity were applied for both pairs, while plasticity could occur either both in the body and the brain or only in the brain. Fig 1 shows the experimental setup along with all the combinations tested. Additionally, a *baseline* experiment was included in which the organisms had to perform a single behaviour for 30 seconds: locomote right. The baseline represents a case in which the organisms evolve in a focal environment, as opposed to having to cope with multiple environmental conditions. Evidently, there was no plasticity in the baseline.

**Fig 1.**
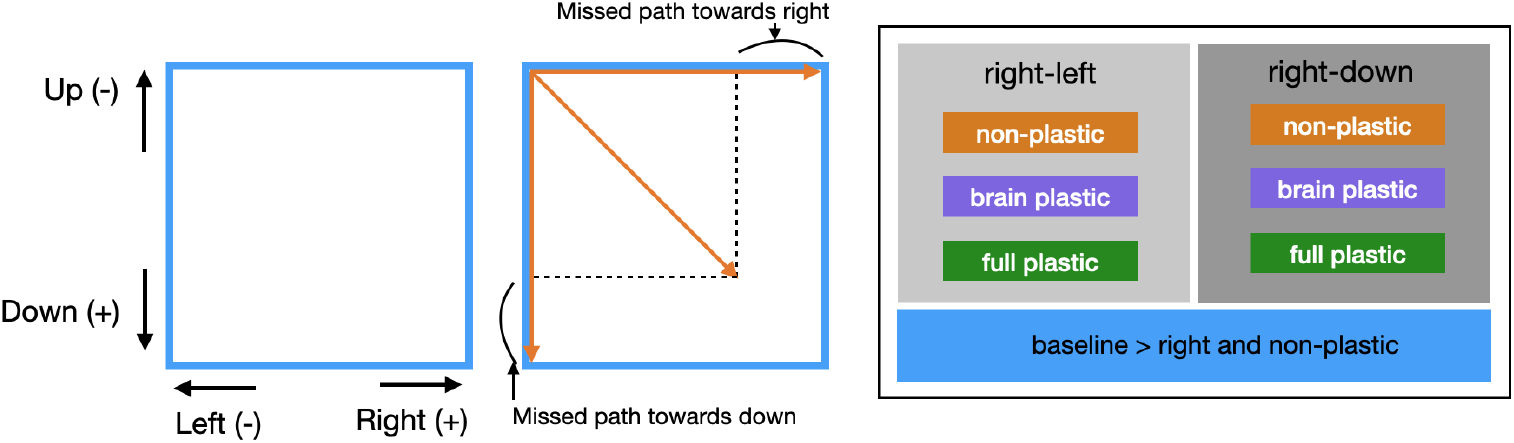
Experimental setup. (left) Directions within the environment. (middle) Examples of trajectories followed by an organism. (right) The different experiments in the setup. *Non-plastic* means no plasticity at all, *brain plastic* means plasticity in the brain phenotype, and *full plastic* means plasticity in the full (body and brain) phenotype.

The design choices made when forming the pairs of desired directions in the experimental setup aimed at composing extreme cases. The objective was to clearly demonstrate both the potential and limitations of the studied approaches. The pair *right-left* requires mutually exclusive behaviours, meaning that behaviour that succeeds at locomoting to one direction fails inversely proportionally at locomoting to the other: by displacing maximally to the right, an organism thus succeeds maximally at displacing to the right but fails maximally at displacing to the left, and vice-versa. This pair illustrates environmental conditions in which behavioural success is either impossible or highly unlikely because the nature of the required behaviours (or phenotypic traits) is too divergent. On the other hand, the pair *right-down* contains directions that can at least partially benefit from each other. This benefit can occur because behaviour that follows a diagonal between right and down results in displacement both towards the right and towards down.

The relevance of these extreme cases has to do with exploring the dynamics of phenotypic plasticity when dealing with a change in the behavioural needs of the organisms. Organisms endowed with plasticity can adapt their phenotype to cope with environmental changes. Therefore, the need to carry out two contradictory behaviours does not necessarily pose a dramatic challenge, even if the behaviours are mutually exclusive. Regarding non-plastic organisms, since their phenotypes can not adapt in response to changes, the need for contradictory behaviours is bounded to result in somewhat poorer behavior in at least one of the conditions.

The success criterion (fitness) for assessing behaviour measured the speed (cm/s) towards the desired direction. Note that the term fitness here refers to a measure of quality calculated *apriori* [31], as opposed to *aposteri* like in biology. It is used to select organisms for reproduction and survival. The final fitness function that consolidates performance amongst the two directions counts the number of Pareto-dominated solutions using the two fitness values: speed in direction A and speed in direction B.

## Results and Discussion

An analysis is presented comparing the ability of each method to produce organisms capable of coping with behavioural changes over the course of their lifetime. The results of these methods are compared to the baseline, to understand if the methods are able to produce similar traits to what they would if only one behaviour was required (focal environment). A video displaying several organisms behaving in the environment is available online https://www.youtube.com/watch?v=EB3-mBoopFY&t=4s.

### Effects of plasticity on behavior

As discussed in the previous section, while for the *right-left* pair phenotypic plasticity would be advantageous, the same can not be presumed about the *right-down* pair. For example, a perfect behavioural repertoire would consist of moving in a straight line towards the left when the expected behaviour is going left and moving in a straight line towards down when the expected behaviour is going down. The lines resulting from these two trajectories form an angle of 90°. Hence, if an organism were to attain a perfect balance between displacing right and down, then this would require moving in a 45°diagonal between the two trajectories. Note that, as depicted in Fig 1 (middle box), this diagonal would leave more than a fourth of the trajectory uncovered (in both directions). However, it is not evident if this ‘loss’ would be sufficient to create pressure towards phenotypic plasticity. For instance, the potential gain of approaching the full trajectory might not pay off when contrasted with the potential genetic costs.

Interestingly, the results corroborate this intuition about the payoff, as demonstrated in Fig 2. Plastic organisms produced a much better performance in both directions of the pair *right-left* than non-plastic organisms, demonstrating that the regulatory system is functional. Considering the fitness function, the fact that the non-plastic organisms on average can barely locomote is not unexpected: solutions are random in the beginning, insofar as there is an equal probability that they displace a little bit to the correct or incorrect direction so that their displacement is close to zero on average; because the behaviours involved are mutually exclusive, the non-plastic organisms never dominate each other, and therefore the zero-average region is never abandoned. Note that although the curves of the non-plastic organisms are completely flat at zero, this does not mean that the organisms from that population did never locomote: this flatness results from aggregating values using medians.

**Fig 2.**
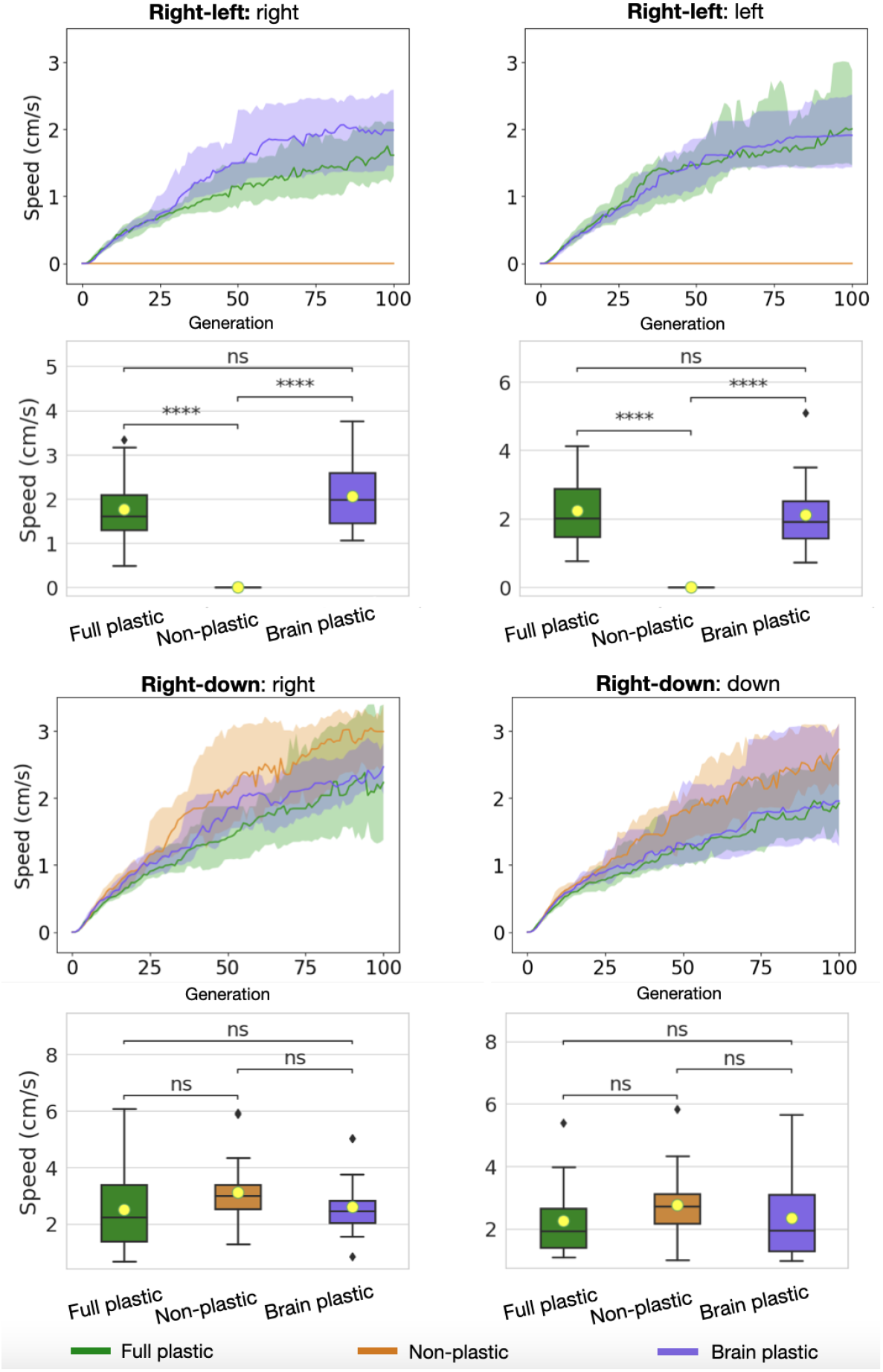
Fitness comparison. *Speed* is averaged within the population for each generation using medians and then averaged again amongst the experiment repetitions using quartiles (lines are the medians, and shades are Q25 and Q75). The boxes compare populations in the final generation using Wilcoxon tests with significance: *ns* : *p* > 0.05; * : 0.01 < *p* <= 0.05; ** : 0.001 < *p* <= 0.01; * * * : 0.0001 < *p* <= 0.001; * * ** : *p* <= 0.0001. The same aggregation and tests apply to the upcoming line and box charts.

In contrast, no significant difference in performance was observed between plastic and non-plastic organisms in any direction of the *right-down* pair. It is relevant to note here that the speed in direction A was comparable to the speed in direction B for all the direction pairs and methods (Fig 5).

These aforementioned behavioural differences are illustrated in Fig 3, which draws the trajectories taken by the organisms when locomoting. As expected, the trajectories of the plastic organisms in the *right-left* pair follow opposite paths when they are required to go either right or left, whereas the non-plastic organisms simply wobble in the middle. Furthermore, the trajectories of the plastic organisms in the *right-down* pair often spread towards the right or down more assertively, according to the due direction, whereas the trajectories of the non-plastic organisms more frequently form a diagonal. Nevertheless, these diagonals appear to roam longer paths than those roamed by the trajectories of the plastic organisms. This visual inspection is aided by measuring the total displacement: non-plastic organisms displace more when summing up *x* and *y* displacement (Fig 4). Such displacement allows the points in space that are achieved by these diagonals to be, on average, as distant from the initial points as the points achieved by the plastic methods.

**Fig 3.**
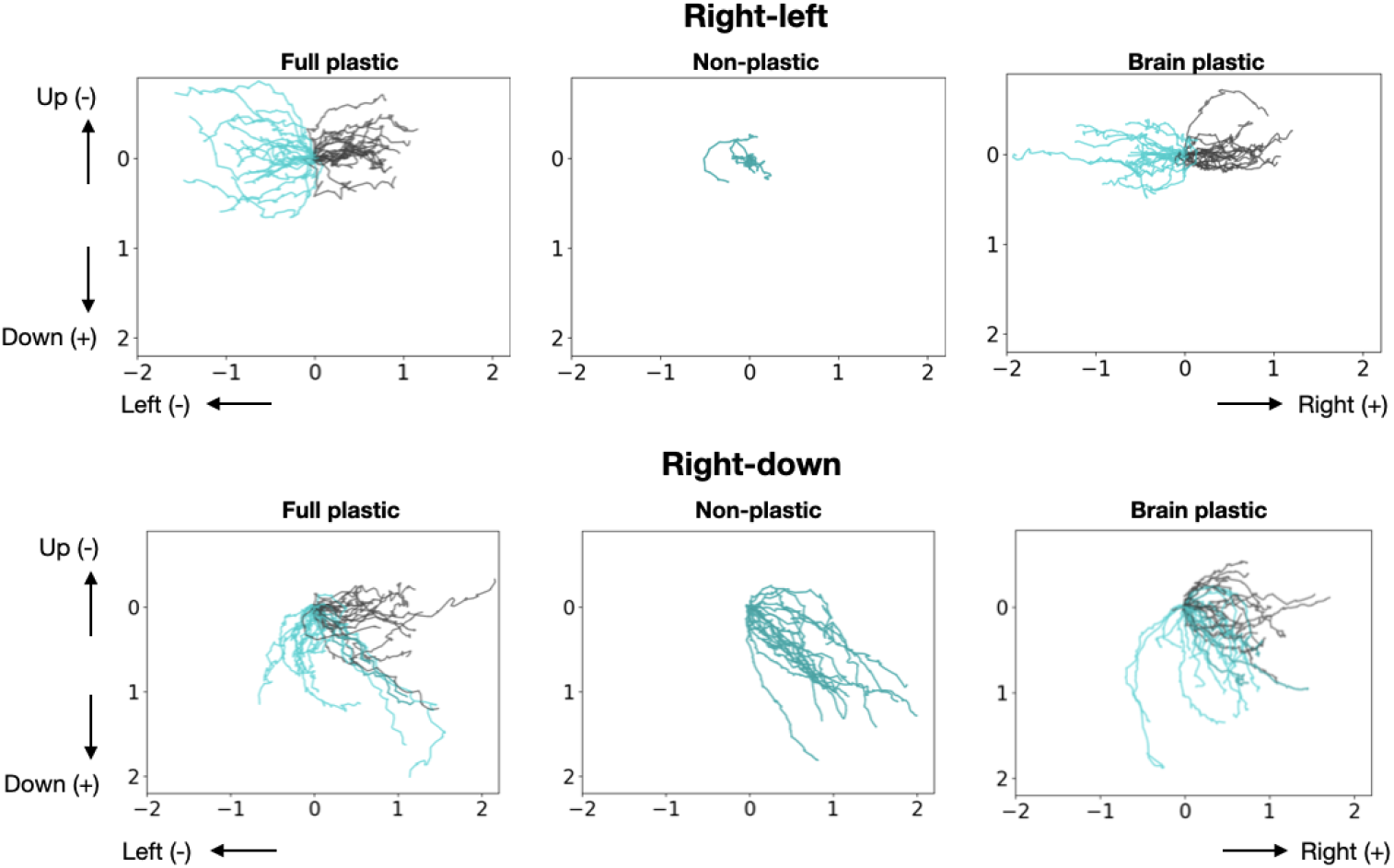
Trajectories of the best organism in the last generation of each experiment. The black and blue lines represent the trajectories of the first behaviour and second behavior of the directions pair, respectively. The trajectories of the experiments were drawn on the same picture, and the individual trajectories of *right-down* can be seen in Figures 1, 2, and 3 of S1 Appendix for better readability.

**Fig 4.**
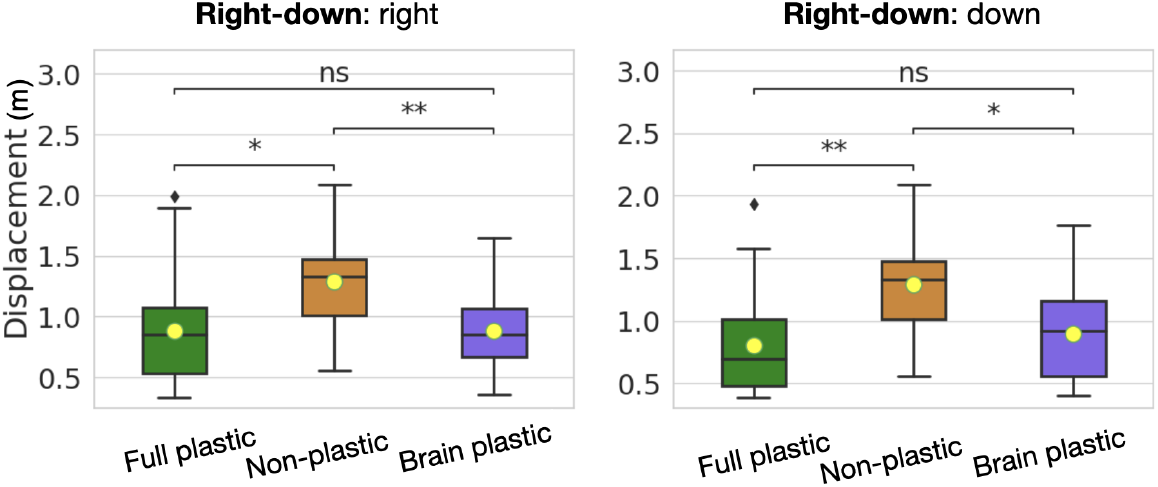
Average total displacement of the organisms in the final generation. Calculated as displacement in the *x* axis plus displacement in the *y* axis.

Finally, the speed of the baseline (focal environment) organisms is much higher than the speed produced by any of the methods (Fig 5). This loss in fitness is expected when comparing the baseline to the non-plastic method because a single phenotype was ‘divided’ between pressures for different behaviours. Nevertheless, such loss was less expected from the plastic methods because they are endowed with the capacity to develop. However, as shall be discussed in the next section, phenotypic plasticity involved costs.

**Fig 5.**
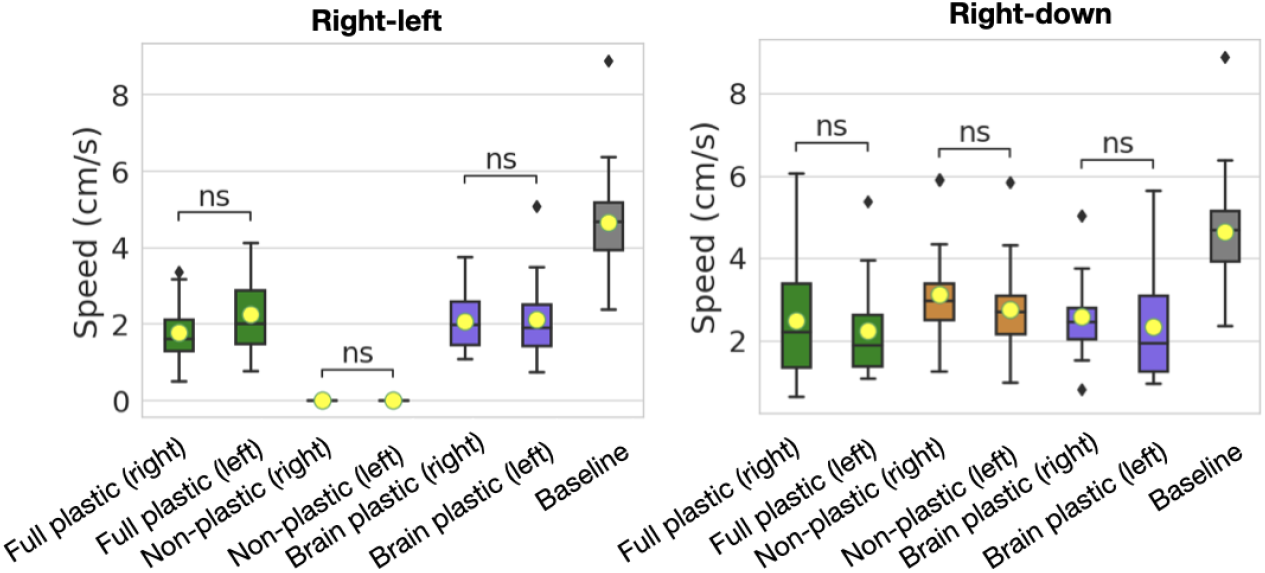
Comparison of speed between the different directions of the pairs of directions as well as in relation to the baseline. Statistical significance markers for comparisons with the baseline boxplot were omitted for the sake of readability. For all boxes, the significance level is between ** and * * **.

### Fitness landscape analysis

The previous analysis demonstrates that beyond being unable to attain the fitness of the organisms from the focal environment, in addition to this, the plastic organisms can not always achieve a superior behavioural repertoire to the non-plastic organisms. To investigate this occurrence, the data were inspected further by analysing the fitness landscape via the following procedure: a search was carried out to explore the space of the possible organisms ‘agnostically’ - the objective of the search was simply sampling the space; the organisms resulting from the search were then measured in regard to body traits; these traits were reduced to two principal components, and the average speed (between the two behaviours) was laid on the landscape within these components. Further details on this procedure can be found in the Methods section.

The procedure produced the landscape illustrations presented in Fig 6. Comparing the three landscapes shows that the *non-plastic* method has fewer, narrower, and less spread peaks (areas of high fitness) than the plastic methods. These observed distributions begin to suggest that the plastic landscapes are more rugged and, as such, more difficult to search. It is important to acknowledge here however, that concluding categorically that the plastic fitness landscapes are more rugged would be an overstatement because we do not know if this is the case for all the possible dimensions, i.e., body and brain traits that were never formulated and therefore not measured. Furthermore, the aggregation applied makes it difficult to interpret the data; for example, speed was averaged using both behaviours as well as the traits in both conditions for the case of body plasticity. Notwithstanding this fact, this examination nevertheless reveals that, at least from these specific dimensions, there is a discernible difference between the studied landscapes. It is thus possible that plasticity influences the ‘complexity’ of the landscape and that this could have played a role in the shortcomings presented by plasticity. This evidence favours the conjecture put forward in the previous section that the potential gain of approaching the full trajectory might not pay off when contrasted with genetic costs. These costs could be reflected as an intensification in the ruggedness as a result of pleiotropy and epistasis.

**Fig 6.**
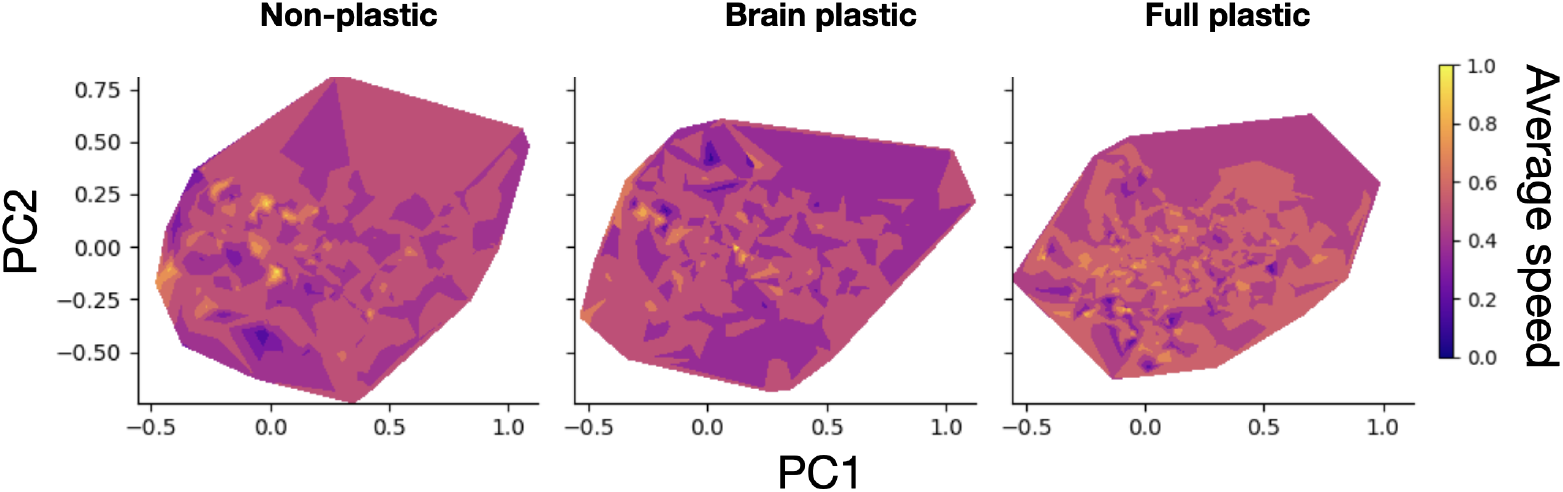
Fitness landscape comparison between methods for the *right-down* pair. Axes represent the components that encode body traits. The components were obtained by Principal Component Analysis of a set containing the data from all three methods. Average speed is the mean of speed in direction A and speed in direction B. The white area refers to values that did not occur.

### Body vs Brain plasticity

Finally, it should be highlighted that endowing organisms with plasticity in both their bodies and brains did not bring about any significant behavioural differences when compared to endowing organisms with brain plasticity alone. The benefits (or the lack thereof) were the same regardless of the extent to which plasticity was allowed. The idea that plasticity is not necessary in the body is further evidenced by comparing the bodies before and after regulating plasticity - the moment when the environment changes and the phenotype develops. The curves of Body Changes in Fig 7 show that as the evolutionary search progresses, the occurrence of plastic changes in the body increasingly reduces so that the final populations are composed mostly of organisms that have the same body in both conditions despite having the capacity for body plasticity. Predictably, this occurs more predominantly in the pair *right-down*, which is expected to benefit less from plasticity.

**Fig 7.**
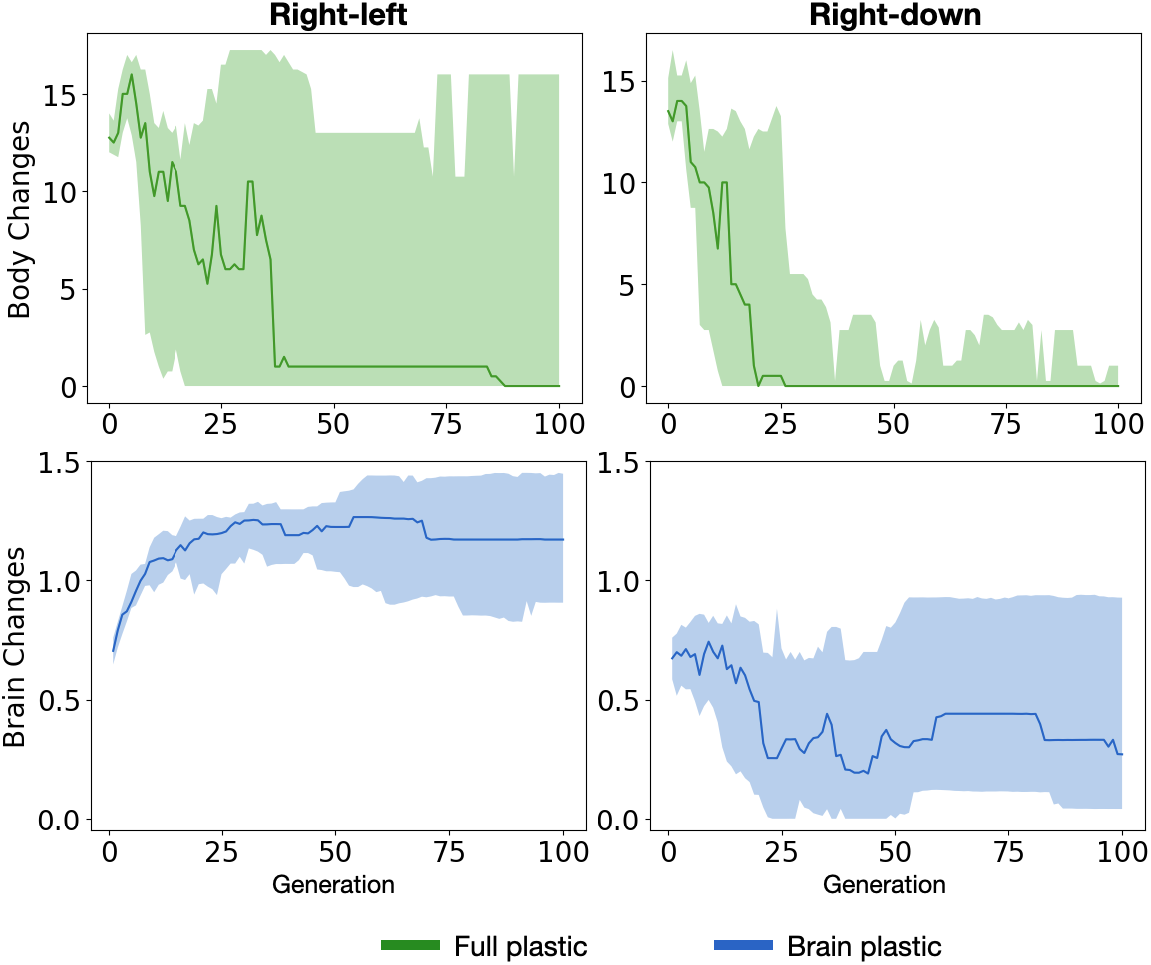
Phenotypic changes. *Body Changes* measures the differences in module configuration between the two possible bodies (one in each condition) of organisms - comparisons were applied only for the *full plastic* method as it is the only one with body plasticity. *Brain Changes* measures the differences in synaptic connectivity between the two possible brains of organisms - comparisons were applied only for the *brain plastic* method and not for the *full plastic* method because the latter allows for changes in brain morphology, which would make comparisons difficult. Details on the metrics are available in the Methods section.

This reduction does not necessarily mean a selective pressure to reduce body plasticity *per se*, but rather could derive from pressure for specific body traits that are optimal for locomotion: this can be observed through analysing how these traits change across the evolutionary period. For example, in both directions of each directional pair, there is pressure for the proportion of joints in the body to increase (Fig 8). Note also here that the final average joint proportion does not differ when compared to both the different methods and the baseline. While there is a difference between non-plastic and plastic in the *right-left* pair, this difference is not relevant, insofar as the non-plastic search was (understandably) unable to leave the random initial region, as previously discussed. This was the case for almost all body trait comparisons: either no difference or no relevant difference was found. The charts showing both the progression and distribution of other body traits are available in S2 Extra analysis. The body traits were measured using descriptors proposed in previous research [32].

**Fig 8.**
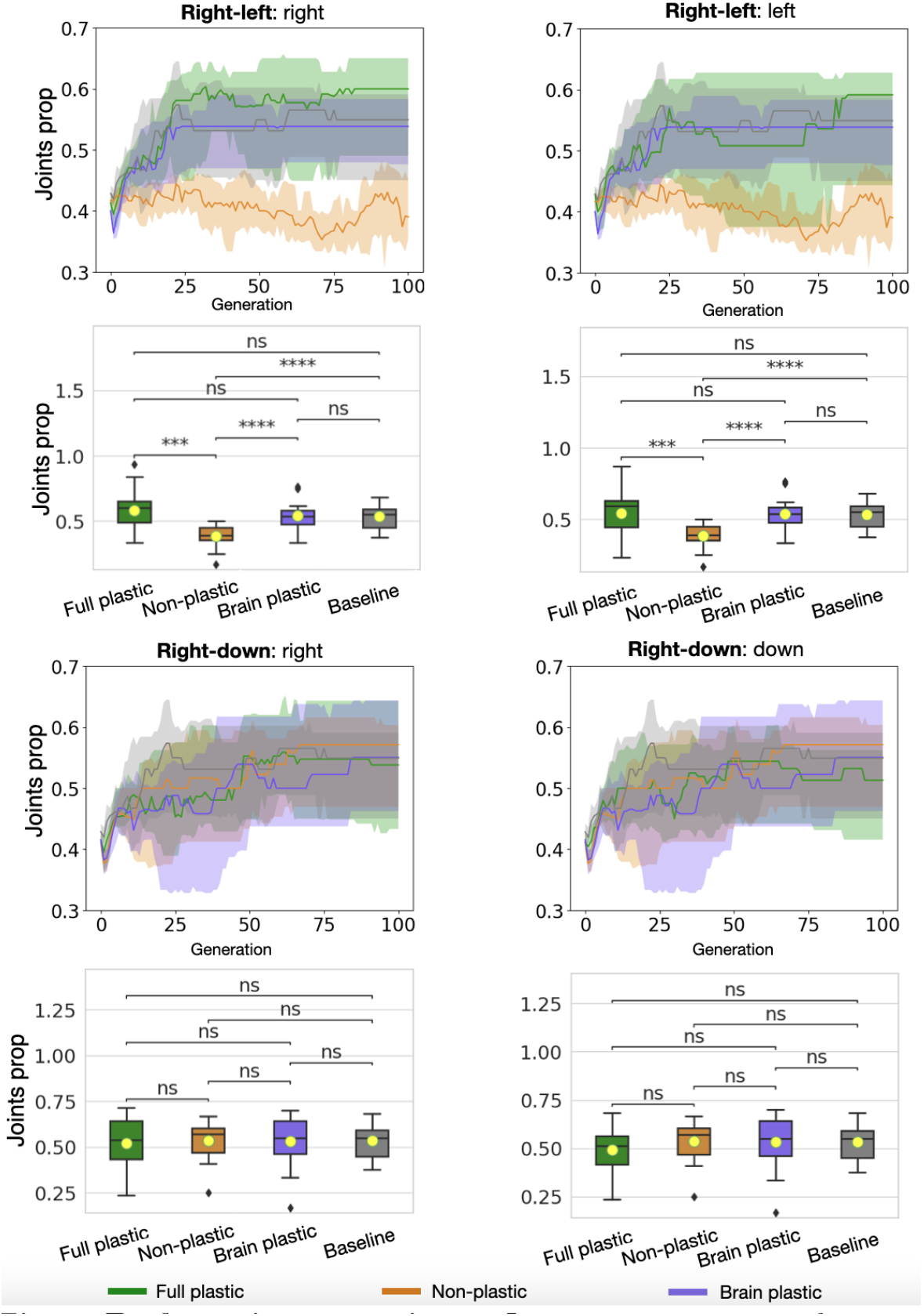
Body trait comparison. *Joints prop* measures the proportion of motor joints relative to the number of modules in the body.

The fact that both emergent phenotypic traits and speed are comparable between the pairs of desired directions suggests that these traits are sufficiently good at the task of directed locomotion irrespective of the direction. Moreover, it must be highlighted that the predominance of certain traits is not due to any obvious developmental bias, insofar as the resulting organisms appear very diverse, as can be discerned in Fig 4 of S1 Appendix.

Obtaining the same traits for both behaviours is unsurprising given that changing the direction does not really change the task. However, the occurrence of this same effect for all methods was not granted, because the challenge was composed of a pair of directions that required distinct behaviour: it is not simply directed locomotion but includes the need for behavioural change during the lifetime of the organism. In this scenario, it is not unreasonable to imagine that certain body traits are favoured to cope with directional changes when there is only brain plasticity, e.g., bodies that do not have limbs in the way of the desired directions. Interestingly however, this favouring turned out to not be the case: brain plasticity itself was able to cope with controlling limbs according to the behavioral needs of each direction.

Conversely, the occurrences of plastic changes in the brain increase across the evolutionary search for the *right-left* pair (Fig 7), which is the pair that benefited from brain plasticity. Regarding the *right-down* pair, while there is no pressure to increase brain changes, but changes are not pushed to zero like in the case of body changes.

## Conclusion

This study investigated the environmentally regulated phenotypic plasticity of evolvable digital organisms. These organisms had to cope with two different environmental conditions (behavioural needs) during the span of their lifetime. Their traits and behaviours were compared to organisms that did not possess phenotypic plasticity. In addition, both these plastic and non-plastic organisms were compared to (non-plastic) organisms that had to cope only with a focal environment.

The results demonstrated a case in which plasticity was beneficial in comparison to non-plastic organisms and a case in which it did not make a difference. In both these cases, plasticity was unable to achieve fitness comparable to the fitness of the genotypes evolved in the focal environment. Moreover, the body traits of the plastic organisms presented no difference to the traits of organisms that evolved only in the focal environment: plasticity thus did not present limits. It is important to acknowledge here that this observation is limited by the current challenges associated with measuring phenotypic traits. In both biology and artificial life, organisms are composed of diverse phenotypic traits from which a multitude of aspects might be derived. Hence, although this study has used multiple metrics to investigate body traits, they might not necessarily capture every aspect of a particular body morphology. Furthermore, metrics to describe brain structure remain missing. Therefore, there can be differences between the plastic organisms and organisms from the focal environment that are not reflected in the current metrics. If these differences exist, they could be interpreted as limits deriving from genetic costs. Similar links between limits and costs have been discussed before [13].

The original hypothesis put forward here was that despite the potential benefits of plasticity for benefits, these benefits might be undermined by plasticity-related genetic costs. The results support this hypothesis, insofar as plastic organisms can attain the traits of organisms from the focal environment, but there are deficits in fitness. These deficits happen in both of the tested scenarios: *right-left*, where the desired behaviours are mutually exclusive, and *right-down*, in which the behaviours are only somewhat contradictory. The two setups are complementary: results from *right-down* alone could leave one wondering if fitness deficits are not simply a result of having to cope with multiple conditions while the regulatory system is idle. However, the *right-left* pair demonstrates a case in which the lack of plasticity results in much more severe fitness deficits. This clarifies that plasticity is functional but leads to deficits and that depending on the expected gain, there might be no benefit in plasticity.

The only plasticity-related costs possible within the utilized artificial life system were genetic costs, this way removing confounding effects that could be generated by other classes of costs, e.g., production costs. Therefore, it is assumed that these deficits derive from genetic costs. This is further evidenced by the landscape analysis, which demonstrated that the landscape is more rugged when plasticity is allowed. When trying to translate the related concepts - the dynamics of the costs of pleiotropy and epistasis - to the context of the current artificial life setup, it is nontrivial to establish a clear map between comparable phenomena. Nevertheless, one might go as far as to hypothesise that the gene interactions and gene reuse between what codes for regulatory capacities and what codes for traits might explain the ruggedness of the inspected landscapes. For example, a particular gene might contribute to both the ability to produce a specific trait and to regulate the expression of this (or another) trait: the first case is about the building blocks of the trait whereas the second pertains to how to put these blocks together. In this scenario, it is possible that a mutation of this gene could result in increasing the quality of a trait, while also reducing the precision of its expression - a better trait at the wrong place. If this is true, then it could mean that mutations in genes that code for regulatory capacities cause frequent and severe ‘shifting’ of phenotypic traits that are dependent on this gene. This hypothetical shifting in dynamics could explain why the plastic landscapes are more rugged than the non-plastic ones: if important traits share genetic material with crucial regulatory capacities, then there might be highly specific combinations of (good) traits that can be produced without disrupting regulation. Such dynamics are comparable to the well-known challenge in ALife of jointly evolving body and brain as opposed to evolving only the brain.

To conclude, this investigation has shed light on how the genetic costs of phenotypic plasticity can occur as a phenomenon in an artificial life system. Although the simplifications of the system make it difficult to evaluate the biological implications of the results, this work contributes to the discussion regarding genetic costs. Finally, the presented results inspired new questions: Under which conditions would the benefits of plasticity compensate for its associated costs? What is the impact of introducing additional costs, e.g., production costs, like energy and matter? How divergent should desired behaviours be, so that brain plasticity is insufficient, thus creating pressure for body plasticity? Under which conditions would one encounter the limits of plasticity? Each of these questions constitutes a potential direction for future research in the field.

## Methods

The code to reproduce all of the experiments and analysis conducted in this research is available on GitHub https://github.com/karinemiras/revolve2/commit/43a6d3915ad87dc0bd192b6db40c8eab4a3f9bc0. The data generated in the experiments are available on Kaggle https://www.kaggle.com/datasets/karinemiras/ploscbio-23-miras.

### Genotype, phenotype, and plasticity

The artificial organisms studied in this work are digital modular robots with a body and brain. The types of modules that can research a body are shown in Fig 9, and the body morphology is optimised via evolution. The brains (controller) of the organisms are Central Pattern Generators (CPGs). “CPGs are neural networks that can produce rhythmic patterned outputs without rhythmic sensory or central input” [33]. While different types of CPGs exist, the current system uses Ordinary Differential Equations (ODEs), i.e., differential CPG [34]. Each controller comprises one or more differential oscillators, which are pairs of neurons that recurrently connect to each other. In addition, neighbour oscillators also recurrently connect to each other as well.

**Fig 9.**
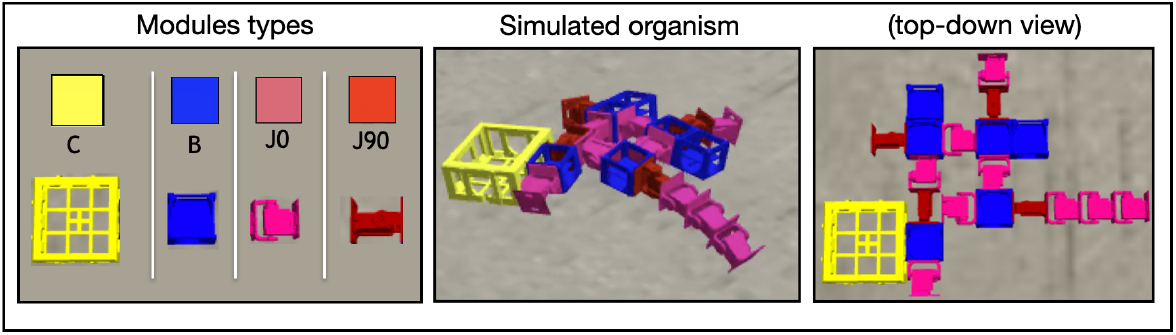
Body morphology. The frame on the left shows the possible modules [35] of which the body of a robot organism. C is the core component, which carries the controller electronic board. B is a structural brick. J is a joint with a servo motor - 0 and 90 refer to the degrees of rotation of the joint in relation to its parent module. The core component and bricks have four lateral slots for attachment. The joints have only two lateral (and opposite) attachment slots. Any module can be attached to any other module through its slots. The modules allow for attachment on their laterals but not on the top or bottom.

The CPG architecture of an organism is closely coupled with its body morphology: there is one oscillator for each servo motor. In this way, the controller of an organism consists of one or more differential oscillators, while each of these oscillators is comprised of a pair of interconnected neurons. For example, the pair of neurons *O*3 and *N*3 forms an oscillator for the motor localised at the point (*x* = 3, *y* = 2) of the grid (Fig 11). These coupled neurons generate oscillatory patterns by calculating their changes of activation at a time step *t* using the following differential equations:

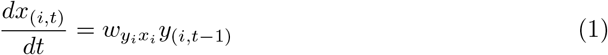

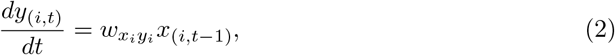

where *x* and *y* represent neurons *O*3 and *N*3 respectively, and *w* represents the weight of a connection between two neurons, and then by updating the values of these neurons through the following expressions:

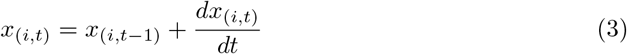

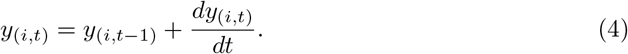

In each time step, the values of the *N* neurons are clipped between −1 and 1 and then sent to the motors.

Compositional Pattern Producing Networks (CPPNs) [36,37] are neural networks that generate structures. A structure is generated by inputting the network with a context related to the structure, such as, for example, coordinates, and then using the outputs of the network to define the building blocks of this structure. The genotypes of the organisms are represented here using CPPNs: one CPPN for the body and a separate one for the brain. The mapping of the genotype into the phenotype is described in Fig 10 and Fig 11 for the body and brain respectively. Importantly, the *environmental regulation* of *phenotypic plasticity* is applied by including additional inputs in the CPPN when querying the genotype to construct the structures. These inputs act as states of that environment that regulate the expression of genetic material into the phenotype (for both body and brain). Generally speaking, such inputs might be composed of any internal or external environmental states (factors) that could be sensed or perceived by the organism. In the current setup, this is provided as a signal that comes from a ‘central system.’ The signal has a value of 1 one when behaviour A is desired, and a value of −1 when behavior B is desired.

**Fig 10.**
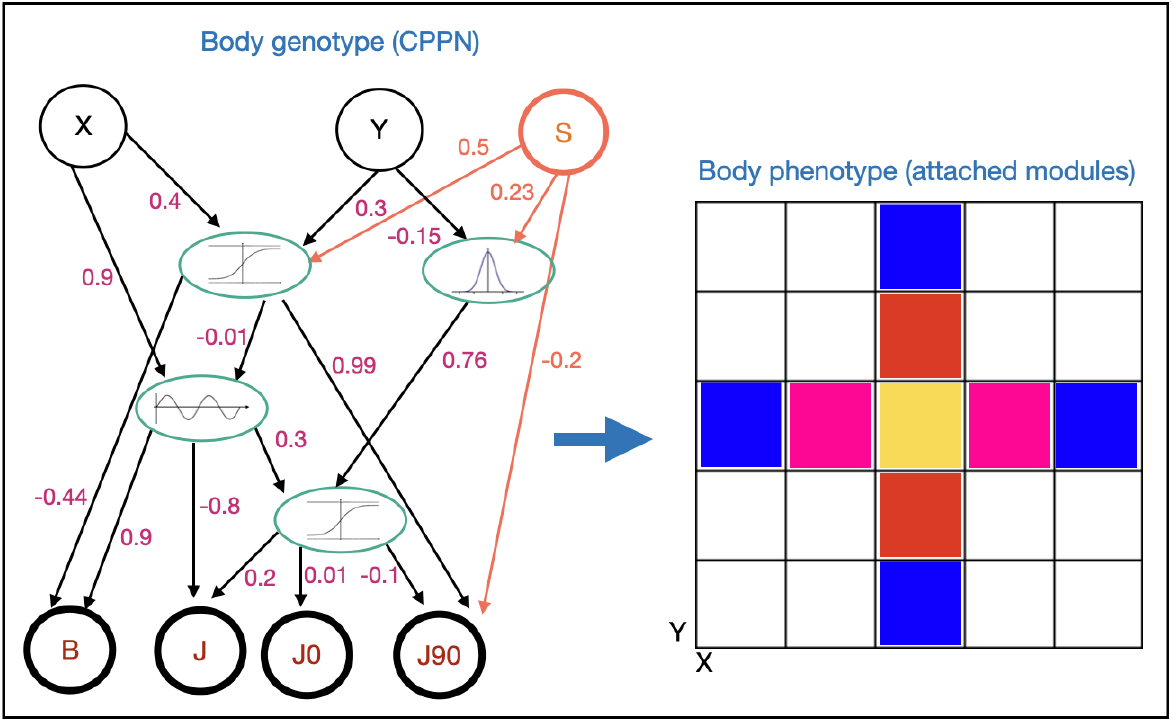
Mapping the body genotype into the body phenotype. A grid with a radius of *r* = 15 points around one central point is defined for the body; the core component is placed in the central point; queries are made to decide which type of module (rotated or not) should be placed in the points (*r* = 2 in the example); querying means providing the *x* and *y* coordinates of a point in the grid as input to the body CPPN and then using the output of the CPPN to decide which type of module should be placed in the point - the neurons with the highest value win the decision of using one module or another and also its rotation; a number of *q* = 30 queries is applied; all points that could be occupied using the available attachment slots have an equal chance of being randomly selected to be queried; as new modules are added and attached, new slots become available; if a query tries to place a module in a point already occupied because of a neighbouring slot, or in a point outside of the grid (marginal slots), then this module is not expressed in the body. Input *S* is a state from the environment used to regulate phenotypic plasticity.

**Fig 11.**
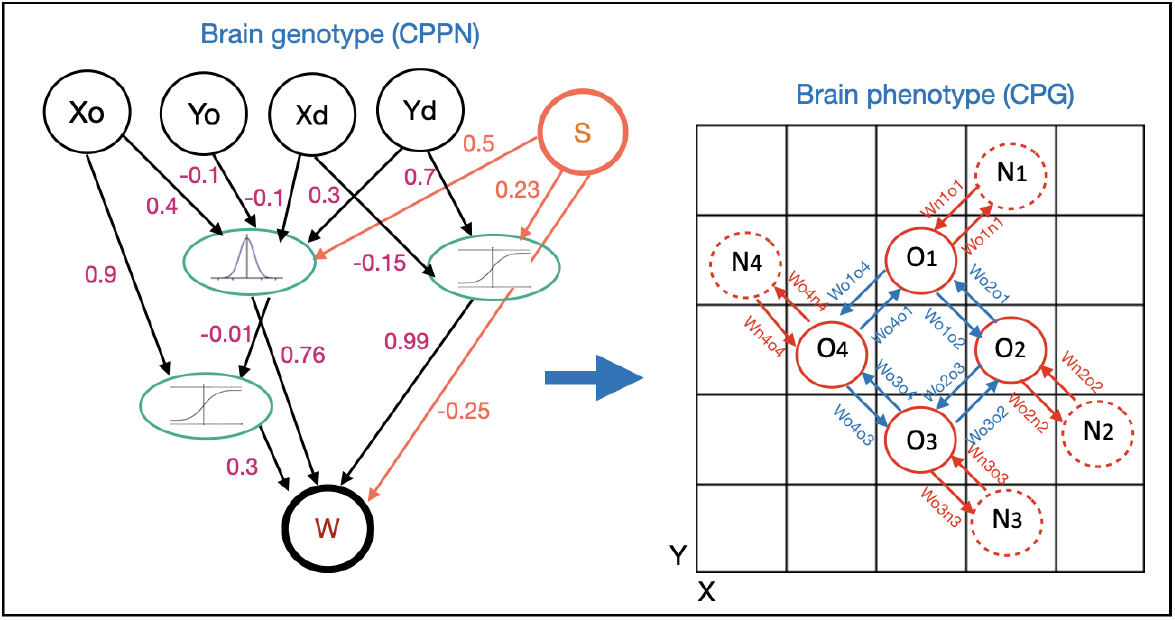
Mapping the brain genotype into the brain phenotype. The brain CPPN is utilized to query the weights of connections between oscillator neurons. The coupled neurons of an oscillator are referred to as (*O,N*). The querying is carried out in two stages. First: the connection within each oscillator is queried by inputting the coordinates of the module to which the oscillator belongs; this value is then used for the connection from *O* to *N*, while its inverse is used for the connection from *N* to *O*. Given that the connections are local, both origin and destination coordinates are the same, that is, (*xo* = *xd, yo* = *yd*). Second: the connections between immediate neighbor oscillators are queried; since an oscillator is composed of two coupled neurons (*O,N*), one of them is chosen as the reference: *O* (drawn with solid lines); in this instance, the origin and destination coordinates are different, because they connect distinct oscillators. The coupled neurons of differential oscillators are drawn in red, while the connection inter oscillators are indicated in blue. Input *S* is a state from the environment used to regulate phenotypic plasticity.

The genotype of an individual is composed of three components: the brain genotype, the body genotype, and a random seed used in the genotype-phenotype mapping of the body. These seeds are required to guarantee heritability in a context in which there is randomness involved in the mapping. This randomness seeks to tackle or prevent any developmental biases. The seeds are not mutated though, since mutating a random seed would result in an unacceptably low locality, i.e., small changes in the genotype that could result in enourmous changes in the phenotype. Hence, in the initial population, different random seeds are generated and then inherited from then on.

### Evolutionary search and mechanisms

A standard evolutionary computing approach was utilised [31]. In each experiment, a population of virtual organisms evolved for 100 generations. Overlapping generations were utilized with a population size *μ* = 200. In each generation, λ = 200 offspring were produced via replication and mutation. The individuals selected for replication were selected by performing binary tournaments. From the resulting set of *μ* parents plus *λ* offspring, 100 individuals were subsequently selected for the next generation using binary tournaments. The fitness function used for each direction measures the speed (cm/s) towards the due direction. The fitness was calculated by evaluating an organism for *30* seconds. Each organism was evaluated for each one of the desired directions independently, while their fitness values were consolidated using Pareto dominance. In this way, the final fitness function counts the number of dominated individuals using the two fitness values (one from each direction). As one would expect, this consolidation was not necessary for the focal environment experiment (baseline).

### Phenotypic changes

The measures of phenotypic changes (from one condition to the other) were calculated by comparing the difference between the phenotype in one environmental condition with the phenotype in another environmental condition.

- *Body Changes*: overlay the two bodies using the head as a reference; count disjunction cases - modules that exist in one body but not in the other; for the intersecting modules, count how many are of different types; add up these two numbers.
- *Brain Changes*: overlay the two brains (they have the same neurons and connections); calculate the absolute difference between the weights of each connection; average all the differences.

### Procedure for landscape analysis

Using the data generated by the studied methods to analyse the characteristics of the landscapes would be unsuitable because these data are biased towards the difficulty of the landscape itself, that is, rugged landscapes might lead to exploring limited areas of the space when using an objective-oriented search. Therefore, additional experiments were conducted to explore the search space ‘agnostically’. This was achieved through Novelty Search (NS) [38], which rather than striving to maximise fitness, instead explores the space to maximise fitness diversity - it discovers solutions that produce fitness values that are maximally different from each other. NS should thus result in the exploration of the space in a non-greedy fashion, which, in turn, allows for a clearer perspective on the landscape.

The fitness function for the NS is defined as *N* = *n*, where *n* is a measure of novelty which is calculated as the average distance to the *k*-nearest neighbours of an organism, for which *k* = 10 and the distance is the Euclidean distance [39] regarding the two desired behaviours: speed in direction A and speed in direction B. The set of neighbors for the comparison is formed by the current population, plus an archive to which 5 organisms of the offspring are randomly added in each generation. The fitness is calculated by evaluating an organism for *30* seconds, precisely as in the other experiments. The evolutionary algorithm utilised is the same as the one described in the previous section, with the exception of the following parameters: population size *μ* = 30, offspring size λ = 30, and number of generations = 100. The experiments were repeated independently 4 times.

To create the landscape dimensions, the following body trait descriptors were utilised: symmetry, proportion, coverage, length of limbs, number of limbs, branching, joints proportion to body size, bricks proportion to body size, and the ratio between rotated and non-rotated joints. Further details about the trait descriptors are available here [32]. All the descriptions range from 0 to 1. Moreover, all measures were averaged between the two environmental conditions to obtain unified values - this includes the trait descriptors and fitness.

Finally, the (averaged) trait descriptors were reduced to two principal components using Principal Component Analysis. The resulting components PC1 and PC2 were placed on axes *x* and *y* of the landscape, respectively, while the (averaged) fitness value was placed on the z-axis.

## Supporting information

**S1 Appendix.** Contains illustrations of the trajectories and body morphology of the organisms and a glossary.

**S2 Extra analysis.** Additional charts generated during the analysis are available on Kaggle https://www.kaggle.com/datasets/karinemiras/ploscbio-23-miras.

## Acknowledgments

I would like to thank my colleague and friend Daan Zeeuwe for his invaluable insight that led to improvements in the genotype-phenotype mapping function that promote diversity.

